# PyReconstruct: A fully opensource, collaborative successor to *Reconstruct*

**DOI:** 10.1101/2025.04.21.649793

**Authors:** Michael A. Chirillo, Julian N. Falco, Michael D. Musslewhite, Larry F. Lindsey, Kristen M. Harris

## Abstract

As the serial section community transitions to volume electron microscopy, tools are needed to balance rapid segmentation efforts with documenting the fine detail of structures that support cell function. New annotation applications should be accessible to users and meet the needs of the neuroscience and connectomics communities while also being useful across other disciplines. Issues not currently addressed by a single, modern annotation application include: 1) built-in curation systems with utilities for expert intervention to provide quality assurance, 2) integrated alignment features that allow for image registration on-the-fly as image flaws are discovered during annotation, 3) simplicity for non-specialists within and beyond the neuroscience community, 5) a system to store experimental meta-data with annotation data in a way that researchers remain masked regarding condition to avoid potential biases, 6) local management of large datasets, 7) fully open-source codebase allowing development of new tools, and more. Here, we present PyReconstruct, a modern successor to the Reconstruct annotation tool. PyReconstruct operates in a field-agnostic manner, runs on all major operating systems, breaks through legacy RAM limitations, features an intuitive and collaborative curation system, and employs a flexible and dynamic approach to image registration. It can be used to analyze, display, and publish experimental or connectomics data. PyReconstruct is suited for generating ground truth to implement in automated segmentation, outcomes of which can be returned to PyReconstruct for proofreading and quality control.

**Significance statement:** In neuroscience, the emerging field of connectomics has produced annotation tools for reconstruction that prioritize circuit connectivity across microns to centimeters and farther. Determining the strength of synapses forming the connections is crucial to understand function and requires quantification of their nanoscale dimensions and subcellular composition. PyReconstruct, successor to the early annotation tool Reconstruct, meets these requirements for synapses and other structures well beyond neuroscience. PyReconstruct lifts many restrictions of legacy Reconstruct and offers a user-friendly interface, integrated curation, dynamic alignment, nanoscale quantification, 3D visualization, and more. Extensive compatibility with third-party software provides access to the expanding tools from the connectomics and imaging communities.

## Introduction

Reconstruct was released more than 20 years ago as one of the first open-source image annotation applications that could be run on a personal computer (1–3), also cite SynapseWeb download cite here. It offers a robust set of annotation features that are user-defined and field-agnostic. Since its release, additional annotation applications have followed, many written with connectomics-level analyses in mind (4–12). These modern applications have prioritized features that allow for rapid imaging and long-range segmentation of cell membranes to identify circuit connectivity (13–16). However, features needed for the more detailed annotation and curation of synapses and subcellular features in the context of circuit-level analyses are downplayed.

Despite the many compelling features offered by the alternatives, Reconstruct continues to be used by a large and diverse group of researchers from a variety of fields. Reconstruct has been used to produce data in hundreds of publications ranging from neuroscience (17–23) to plant biology (24, 25), entomology (26–28), human anatomy (29), herpetology (30), and materials science (31), to name a few. We attribute its widespread use in part to Reconstruct’s simple interface and flexible approach to annotation. Image stacks are imported into the application, and with little effort, researchers can quickly begin annotating serial sections, export data for analysis, and produce publication-quality 3D visualizations, all in a single intuitive application.

Despite significant advances in EM annotation tools, several fundamental challenges persist. Many require considerable programming expertise to access full annotation details and advanced features. This requirement limits accessibility to technical specialists rather than serving the broader research community. Most annotation tools lack robust, integrated systems for quality control and data curation. Many do not accommodate post-hoc realignment of serial sections, making it difficult to correct alignment errors discovered during annotation or analysis. Finally, most tools separate meta-data from image data, forcing researchers to maintain parallel data management systems and complicating long-term data preservation. These limitations can significantly impede annotation workflows and compromise data integrity, particularly in large-scale projects.

Reconstruct has long been overdue for a major overhaul. It only ran natively on Windows machines, datasets were RAM-restricted in size, a complex system of transformations hampered new alignment strategies, and minimal interaction with outside tools restricted its expansion. Limitations like these required users to implement kludgy workarounds that complicated their workflows. Despite these limitations, Reconstruct continues to be used for its simplicity and efficiency.

With these limitations in mind, we have produced a generalizable and streamlined annotation tool in the spirit of Reconstruct that integrates readily into any serial imaging pipeline (Fig. 1). We have named this new system PyReconstruct, a modern, user-friendly successor to what we now refer to as “legacy” Reconstruct. PyReconstruct was written in Python, retaining legacy Reconstruct’s intuitive interface and addressing the shortcomings necessary to transition from serial section to the volume electron microscopy needed for circuit analyses. It features an integrated curation system for maintaining data quality, flexible alignment capabilities, and unified data storage keeping meta-data and image data together. PyReconstruct runs on most operating systems, employs lightweight data structures, implements dynamic alignment, and provides multi-resolution scaling. It lifts RAM restrictions allowing users to scale up to large volumes. PyReconstruct is also backward compatible with data annotated in legacy Reconstruct, giving legacy users the ability to port their data into the modern annotation environment. Importantly, PyReconstruct is fully opensource and users are at liberty to customize the source code to their own use cases.

**Figure 1.**
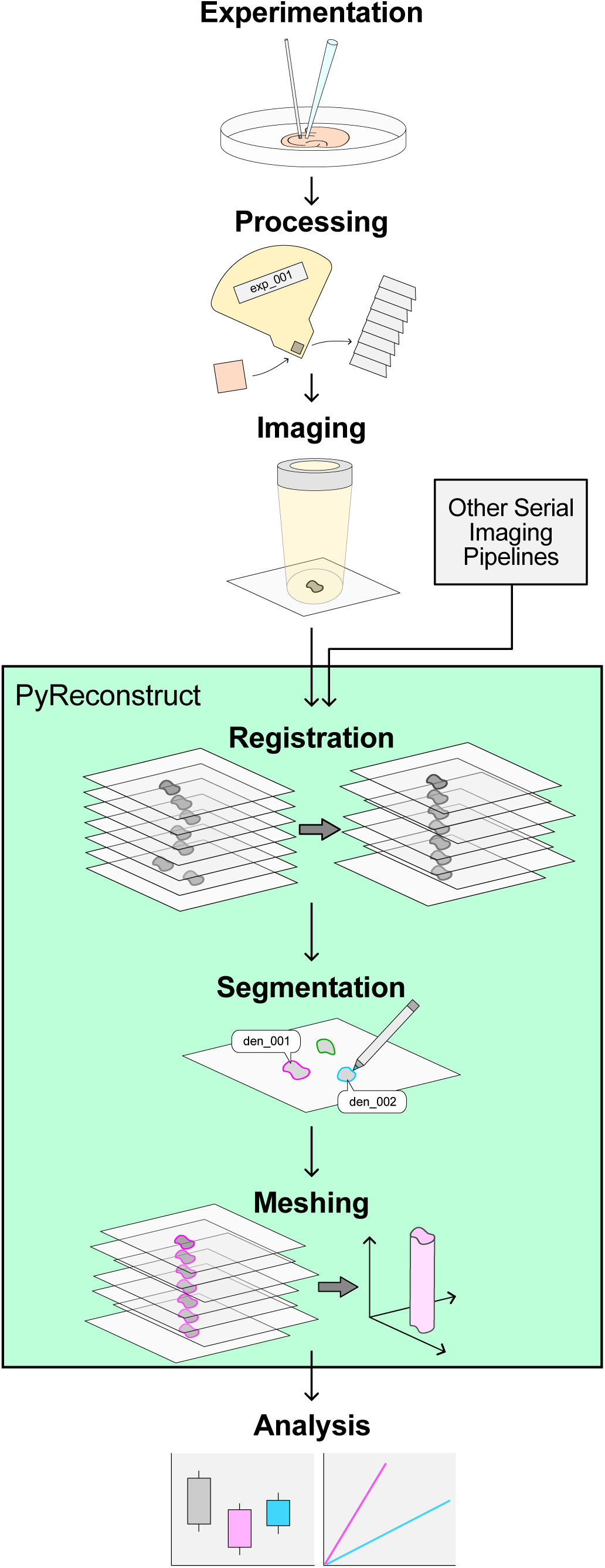
Example serial imaging pipeline from experimentation through data analysis. PyReconstruct encompasses the most time-consuming steps (green box) in pipelines that rely on manual segmentation or proof reading of serial images to produce analyzable data. Here, a serial section EM pipeline is shown. Image registration, segmentation, and contour meshing can be performed in a single application. Images from any source can be used in this pipeline, including series or single sections.

### User experience

We designed the core functionalities and features of PyReconstruct to facilitate working with serial images in an unopinionated manner, to maintain data integrity, and to promote a collaborative workflow. In the sections below, we describe the annotation and segmentation interface, object organization capabilities, curation tools, alignment systems, and data visualization features.

### Segmentation and object organization

PyReconstruct offers an uncomplicated graphical interface for registration, segmentation, and meshing of image data and allows for multiple points of entry and exit to and from third-party software (Fig. 2). By leveraging Python wrappers for openCV (4.8.1), stacks of serial 2D images can be loaded into PyReconstruct in multiple formats (.tiff, .jpeg, .png, etc.). Though PyReconstruct directly imports standard images without conversion, users can also benefit from its support of precomputed volume formats, the advantages of which are outlined below.

**Figure 2.**
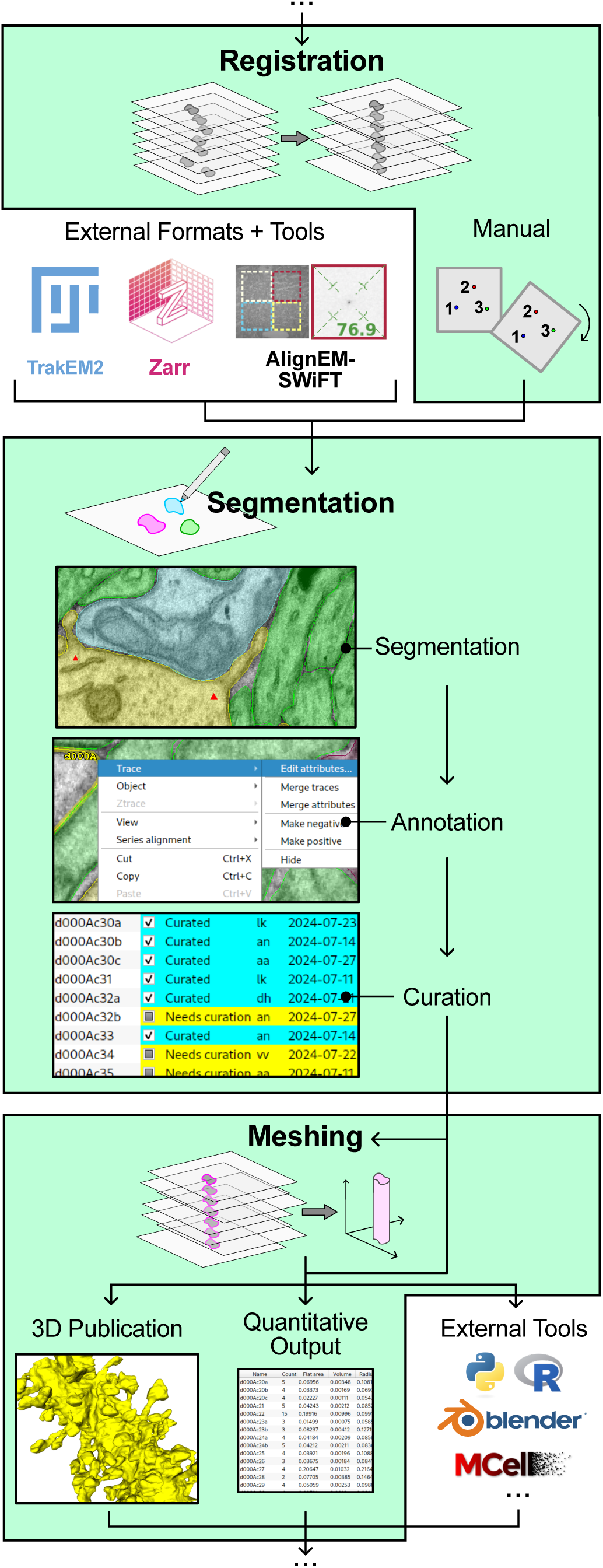
PyReconstruct readily interacts with third-party software. Core PyReconstruct functions are highlighted in green and include native manual registration through the application interface, while also supporting importation of pre-registered images and transformation matrices (6, 32, 33). PyReconstruct stores detailed segmentation data as 2D contours and features a checklist-based curation system for quality assurance. Users can generate, view, and export quantitative data and meshes directly in PyReconstruct to create publication-ready figures or for use externally in programming languages such as Python and R. Exported meshes can be further edited in advanced 3D platforms like Blender for biophysical modeling using programs like CellBlender and Mcell (34–36).

The user’s primary workspace contains a single image from the series (the “section”) displayed in the main window (Fig. 3A). The trace palette, tool bar, section navigator, and brightness/contrast sliders are superimposed over the image and are moveable around the field, such that the user can personalize their workspace as they annotate. Detachable windows display quantitative 2D (the trace list) and 3D annotation data (the object list) as it is being collected in the main window (Fig. 3B). Users are at liberty to choose trace name, color, tag, and appearance. The trace list allows users to view and edit traces on a section and see quantitative measurements, while the object list allows users to view and edit object data and see quantitative measurements in 3D based on the calibrated section thickness (2).

**Figure 3.**
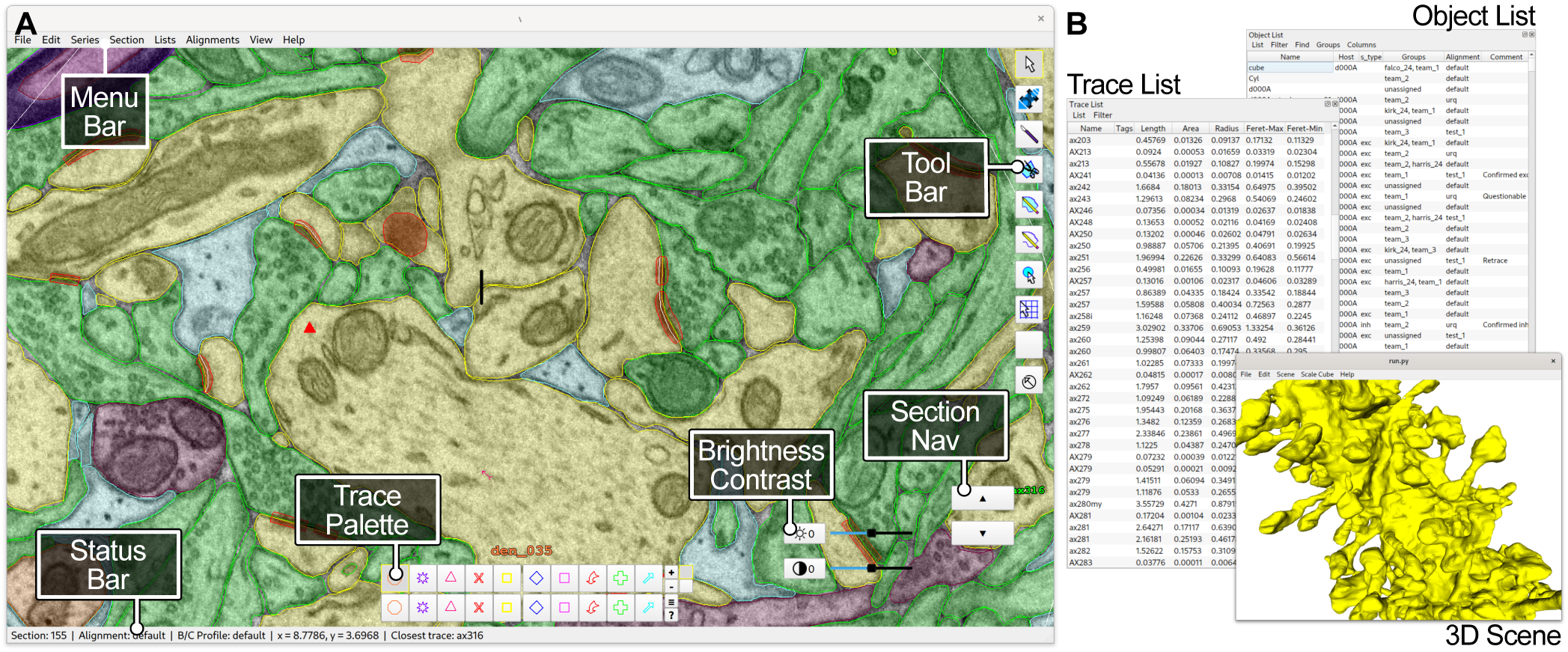
Windowing schema in PyReconstruct. (**A**) The main window contains images and widgets (tools, trace palette, etc.) that can be moved around the frame and toggled on and off, allowing users to customize their primary workspace. (**B**) Users interact with 2D and 3D data as it is being collected through ancillary, detachable windows, such as the trace list, object list, and 3D scene.

In PyReconstruct, 3D “objects” represent collections of 2D “contours” over the image stack that belong to a single feature in the series. An object’s contour on a single section may consist of one or several separate “traces” (Fig. 4A). Segmentation data in PyReconstruct is stored as lists of x and y positions that make up individual traces, and objects can be organized and classified in a number of ways. Hierarchical and nested objects groups can be created by assigning “hosts” that point to other objects. For example, “synapse” may point to “dendrite” and/or “axon”. Users can also create custom categories to assign objects qualitative measures. For example, “synapse” from the above example could also be categorized by “synapse type” as “excitatory” or “inhibitory”. For cases that fall outside host-inhabitant relationships and categorical variables, users can create simple, custom-named groups. Because these classification schemes are customizable, users can define semantically meaningful groups and relationships that are specific to their use-cases.

**Figure 4.**
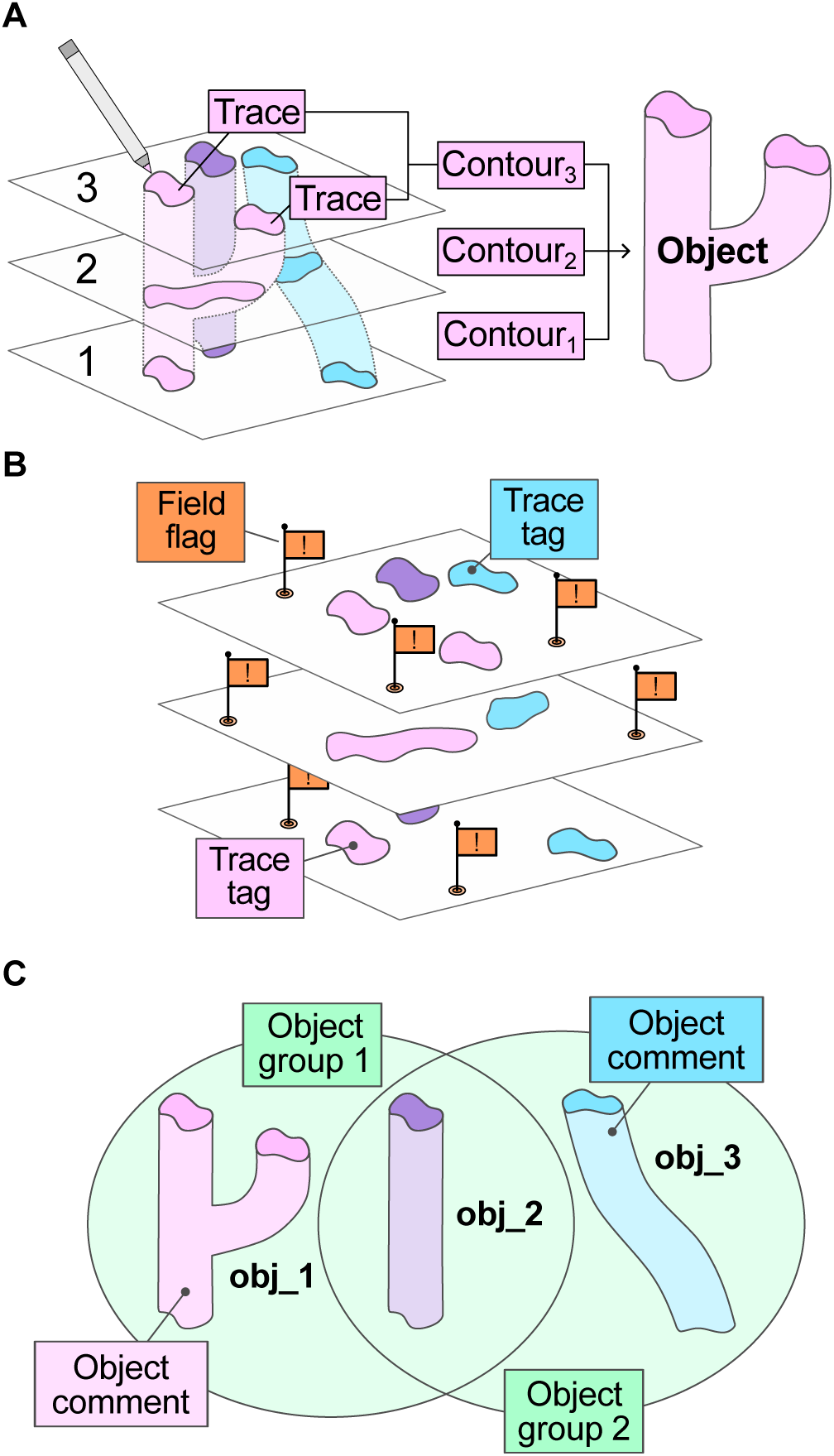
PyReconstruct stores both free-form and hierarchically organized annotation data. **(A**) Object profiles on individual sections are identified and their outlines traced by annotators. An object’s 2D contour consists of all traces on a section belonging to the object. 3D objects are constructed from multiple contours across serial images. (**B**) A trace tagging system allows users to ascribe additional information concerning 2D segmentations. (**C**) Annotations applied to objects include groups and comments, which can be accessed from the object list.

### Rich, free-form annotations and event tracking

PyReconstruct allows for rich annotations that might otherwise be stored externally and risk being separated from the original series. Annotations that provide meta-data and context to series features can be applied to individual traces (tags), entire objects (groups and comments), and points of interest in the field (flags) (Fig. 4B-C). For example, traces denoting object profiles distorted by image artifacts might be tagged to indicate they have been interpolated (or inferred by the user) from traces on adjacent intact sections. Users may flag objects in a series that serve as probands or objects the annotator wishes to revisit later. Objects can be grouped to track their use in specific publications or projects, while comments provide additional explanatory notes. Free-form annotations like these enable users to create and customize their own annotation system based on project needs.

Experimentalists often revisit previously annotated series to test new hypotheses, and keeping track of the history of actions performed in a series may serve to guide subsequent annotation. In the past, this information was stored externally. PyReconstruct provides automation by logging all actions that add new or modify existing annotation data logged. Log entries contain brief descriptions of annotation events, including date, time, user, objects edited, and the section or sections on which the action was performed. Users can view and filter the full log to display entries relevant to one or more objects.

### Collaborative curation for quality control

Annotating large amounts of serial section data is generally performed among teams of annotators and PyReconstruct was developed with this approach in mind. Annotations performed by a team member are subsequently passed to a more expert annotator to be proofread and curated for quality assurance (Fig. 5). To this end, PyReconstruct includes a simple curation system, and users can be assigned curation tasks through the object list (Fig. 6A). Curation history (user, date assigned, completion status, etc.) is tracked with the series, so that project leads can verify if annotation criteria is applied uniformly among team members and correct tracing errors. Curation data and history is readily available and displayed through the object list. Free - form annotations (see Fig. 4B-C) provide further context to aid collaboration among annotation teams. Field flags also include a running commentary that can be used to alert present and future annotators to information about locations in the series (Fig. 6B). Flag meta-data including the creation time, username, and related notes are tracked and stored with series data. A conversation-style comment system allows users to maintain a running discussion for each flagged item. Tags applied to traces on a particular section can be accessed in a trace list (Fig. 6C), which when filtered and sorted, provides a rapid means to identify the status of a trace.

**Figure 5.**
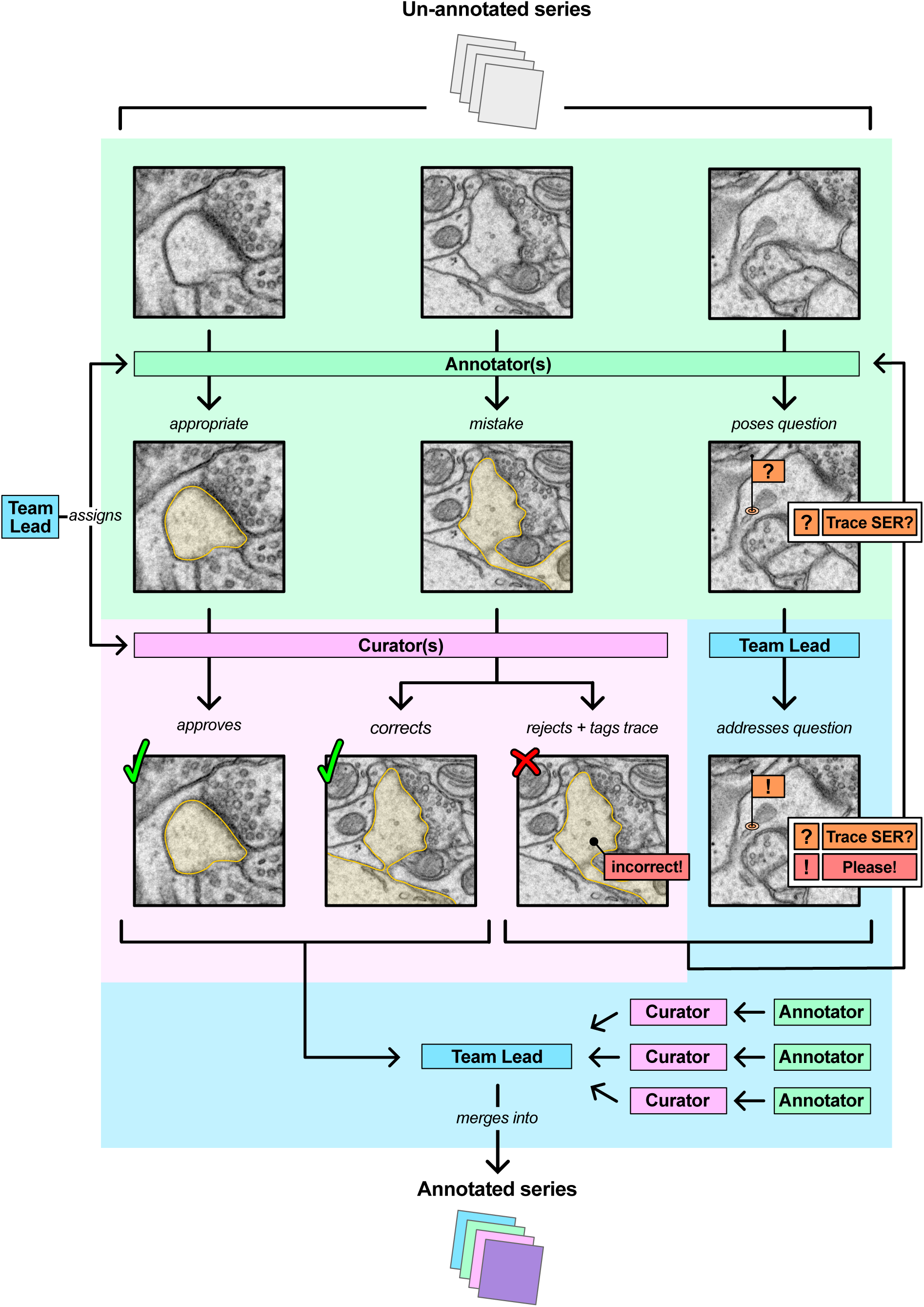
Implementation of a team-based annotation and curation scheme to ensure quality control. A team lead (blue) assigns annotation (green) and curation tasks (pink) to team members working simultaneously on copies of a single series. Curators approve, correct, or reject traces, which can be tagged for review. Annotators can pose questions to team leaders through a system of flags with commentary that can be answered and returned. As annotations are curated, the partially annotated series can be passed back to the team lead. Curated series are merged into a single annotated dataset for analysis. PyReconstruct provides features at multiple steps in this pathway to facilitate curation and quality control.

**Figure 6.**
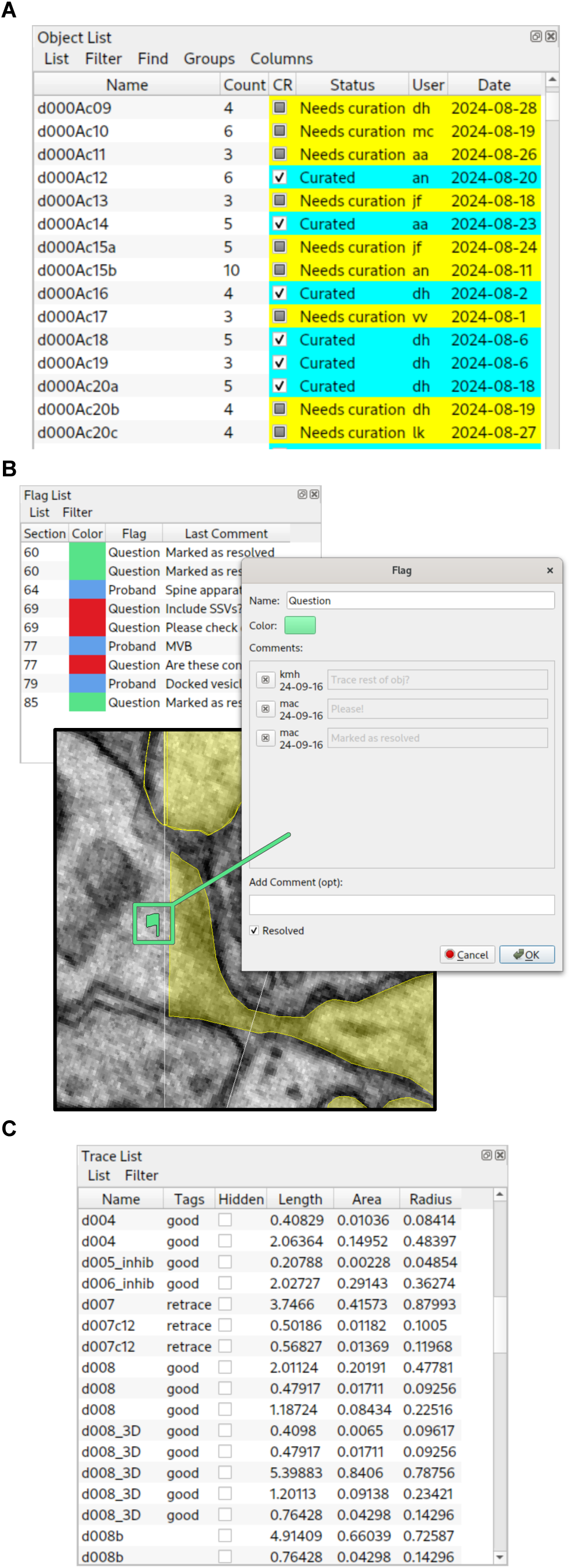
PyReconstruct provides automation for curating segmentation data. (**A**) Tracking of curation tasks is stored with series data in PyReconstruct. Curation tasks and status can be assigned interactively to objects from the object list. Multiple filtering options allow users to view pending and completed curation tasks assigned to them and to others. Here, users were randomized to curation tasks through PyReconstruct’s Python API. (**B**) Flags placed in the field are shown in a flag list, which can be sorted in various ways. Running commentary allows team members to pose questions and discuss issues that arise during annotation. (**C**) Traces can be tagged with information that is accessed from a sortable trace list.

PyReconstruct runs on a local machine and does not currently implement a client-server solution to allow multiple users to work simultaneously on a single series. Trace conflicts might therefore arise when series annotated simultaneously by different users are subsequently merged. For example, two annotators tracing the same object will lead to non-identical, overlapping traces representing the same feature on a section. Merging large, heavily annotated series can result in many duplicate traces that pollute the workspace and lead to quantification errors.

To deal with this problem, a trace import system has been implemented in PyReconstruct that identifies and alerts users to merge conflicts that must be resolved. Conflicting traces are determined based on the Jaccard similarity coefficient, trace history, and annotator preference. Users can determine an overlap threshold above which traces are considered “identical enough”, which gives PyReconstruct leeway to discard all but one overlapping trace. This provides some automation to the tedious tasks of curating multiple traces for a single feature. Remaining conflicts are flagged for resolution, and users can walk through each conflict systematically.

### Dynamic alignments applied on demand

During processing, sectioning, and imaging, planar specimens are subjected to a variety of physical alterations (e.g., compression, shrinkage, heating) and technical complications (e.g., stage and specimen drift, and non-uniform scan and lens characteristics) that introduce linear and non-linear distortion into the final images (37–39). Image distortion can be mitigated through choice of processing protocol (40–43), imaging modality (44, 45), and post-hoc computational processing, but in many cases must ultimately be corrected through image registration. Registration methods are rapidly improving (46–57), and choice in strategy depends on distortion type and personal preference. We therefore sought to implement a flexible and dynamic registration strategy to allow users 1) to import alignments generated externally, 2) to perform simple registration inside PyReconstruct, and 3) to store multiple alignments that can be applied to stacks of images on the fly (Fig. 7).

**Figure 7.**
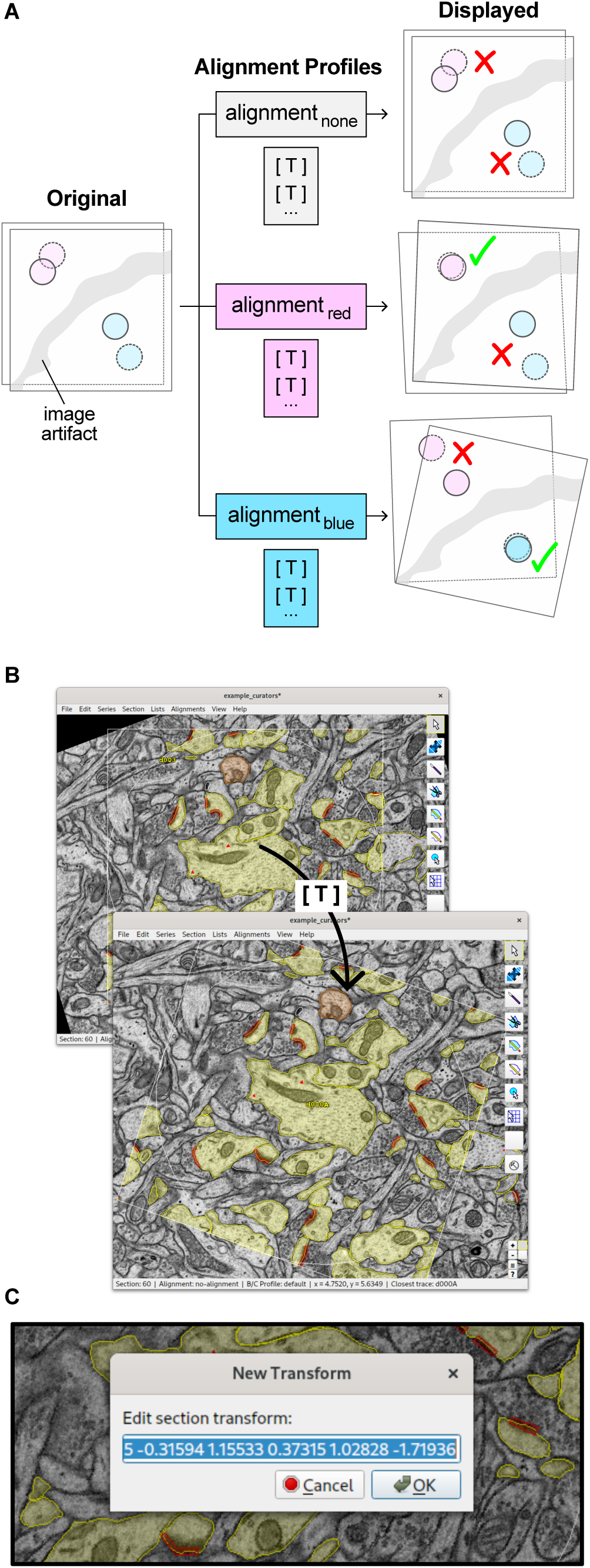
Multiple alignment profiles applied to serial sections on demand. (**A**) PyReconstruct stores image and contour data in their untransformed state. Alignments are represented as lists of affine transformations ([T]), one for each section. Multiple alignments are stored as profiles that can be applied independently on demand. Image artifacts (resulting from tears, folds, etc. in serial sections) often warp serial sections and produce local misalignments. The ability to tweak, create, and store multiple alignments facilitates annotation. (**B**) Alignment profiles can be accessed and modified through the user interface. The current alignment is displayed by name in the main window’s status bar (bottom of main window, see Fig. 3A). (**C**) New alignments can be created in PyReconstruct and imported from external applications or section transformations can be edited directly in the interface, allowing users fine control over series alignments.

An “alignment” in PyReconstruct is represented by a list of affine transformations (Fig. 7A). Each section in the series is associated with a single affine transformation, which is stored as a set of six numbers that correspond to the first two rows of the 2D transformation matrix. This single transformation is applied to both image pixels and contour data of a particular section, which are then displayed in the field when the section is called by the user. In this way, users whose images require affine transformations alone need not work with images whose alignments have been previously “baked in” (i.e., transformations applied directly to stored images).

This strategy offers several advantages. First, the only image data stored with a project are the unaligned images that were initially produced at the microscope and imported into PyReconstruct, dramatically reducing the size of a project. (Users can still of course use third-party software like Fiji, TrakEM2, and SWiFT-IR to import images with baked-in alignments and perform additional local alignments.) Second, stored contour points reflect the x and y position of the points on the unaligned images (not the transformed images), making it possible for users to apply external transformations to those points when exporting contour data from PyReconstruct. Third, users can rapidly tweak a section’s alignment should they note alignment flaws while annotating, which is a common occurrence when working with serial images.

Finally, the simplicity of this strategy means users are at liberty to store multiple alignment profiles specific to an object, region of interest, or context, which can be applied to the series on demand (Fig. 7B). This is especially useful when annotating objects in areas of section deformation, for example, on either side of image artifacts (breaks, folds, tears, etc.), which are frequent in serial section EM. These artifacts would otherwise require users to use third-party image editing software to correct. Thus, a complete history of alignments for each series is stored. Objects in the series can be assigned to specific alignments, ensuring the object’s quantitative measurements and 3D reconstructions are accurate.

PyReconstruct supports simple manual alignment natively and users can interactively translate, rotate, scale, and shear images in the main window. Users can also directly edit a section’s affine transformation through the menu bar (Fig. 7C). Affine transformations can be estimated by identifying fiducial markers in the field, such as cell structures that span multiple sections. Transformations performed on single sections can be applied throughout the remaining sections, allowing for a correction in alignment to be propagated throughout the series. PyReconstruct supports importing alignments from other PyReconstruct projects as well as from external applications, for example, directly from SWiFT-IR project files (33, 56) or in the form of simple .txt files, each line representing an affine transformation.

### Data visualization and analysis

PyReconstruct’s object list represents a serialized view of objects, their measurements, and meta-data, and allows users to export data for analysis and render objects in 3D (Fig. 8). 2D contours are voxelized and represented as NumPy arrays, which are translated through a matrix-to-marching-cubes algorithm provided by the trimesh library (58) to generate watertight triangle meshes for visualization and quantification (Fig. 9A). The trimesh library provides a number of in-place smoothing filters (59, 60) that can be applied to meshes in PyReconstruct. 3D meshes are visualized in a customized 3D scene built on top of the Python scientific visualization library vedo (61). The meshing strategies employed natively in PyReconstruct are modularized, such that users proficient in Python can implement more complex meshing algorithms accessible from the user interface should they choose to do so. 3D meshes can be exported from the object list in several formats (obj, ply, stl, etc.), which can be imported into external 3D modeling software, such as Blender, for further editing.

**Figure 8.**
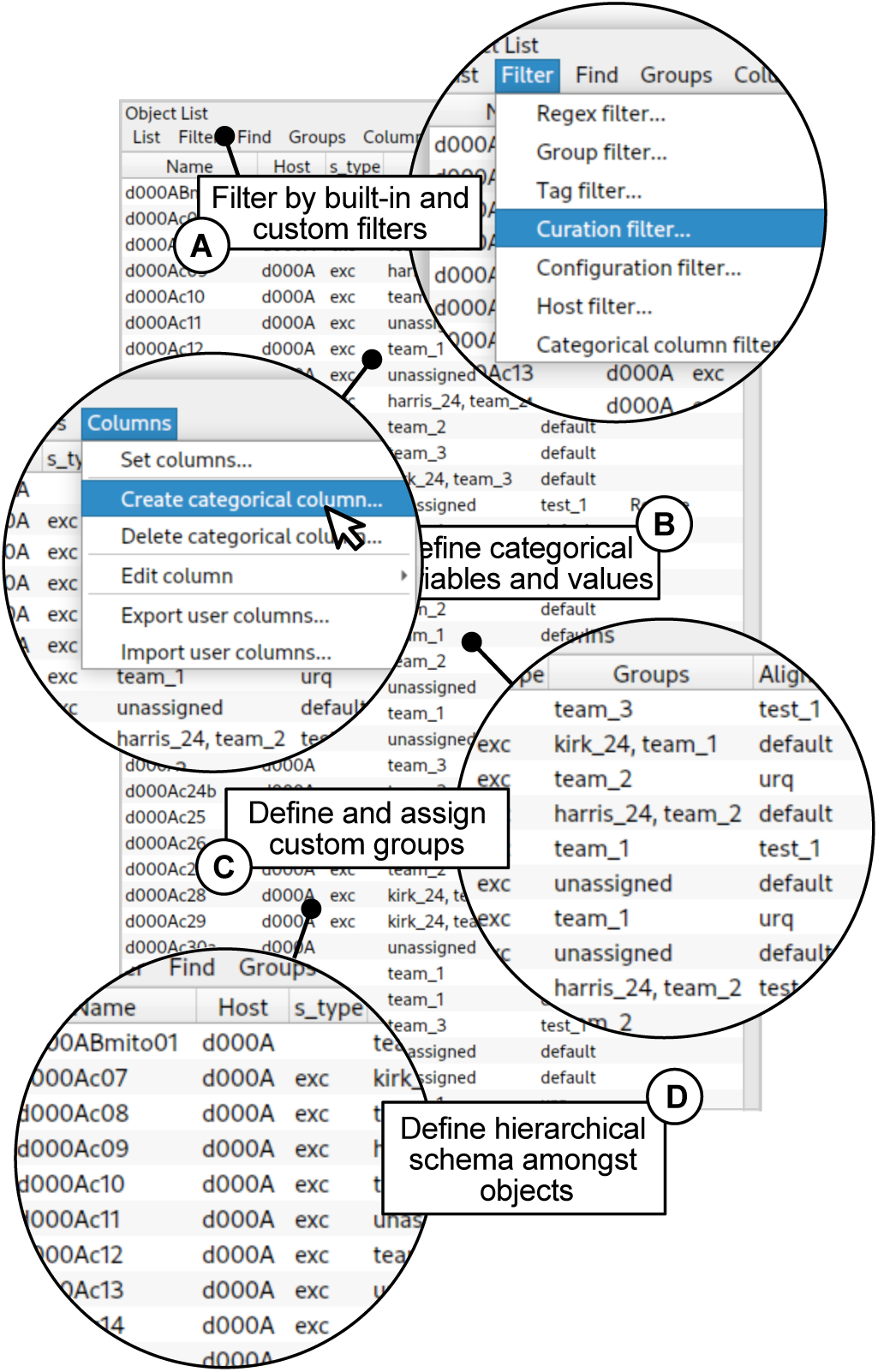
Three-dimensional data is stored in the object list. Data concerning 3D objects constructed from 2D contours is displayed in the object list. Users can sort by built-in and custom filters (**A**), define categorical variables and assign values (**B**), define and assign custom groups (**C**), and define hierarchical schema amongst objects by assigning objects as hosts (**D**).

**Figure 9.**
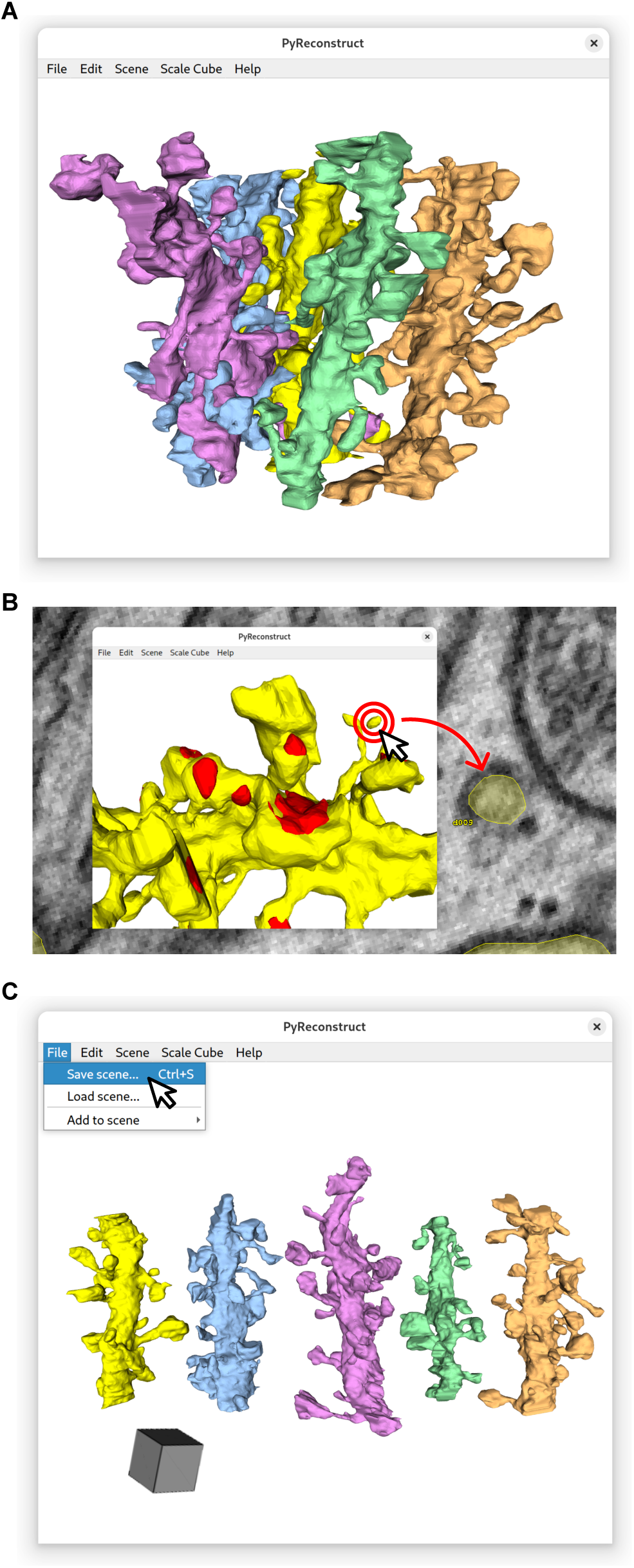
An enhanced 3D scene facilitates detailed curation and stores publication-ready visualizations. (**A**) 3D renderings of objects can be generated from the object list and are displayed in a featureful 3D scene. (**B**) Double-clicking in the 3D scene focuses on the corresponding 2D location in the main window, allowing users to navigate to sections for curation. Here, a disconnected part of an object identified in the 3D scene is rapidly uncovered on the 2D section, allowing users to curate more finely the object’s segmentations. (**C**) The updated 3D scene offers users features that organize objects in an automated fashion. Here, all objects in the 3D scene have been organized along a single axis, providing a visual overview of all meshes. Objects can be moved and rotated interactively and their attributes (color, opacity, etc.) can be altered. Scenes can be exported, saved, and re-loaded at a later time, allowing users to produce publication-ready figures and store scenes with series data.

Segmentation issues are often not apparent when viewing objects as 2D profiles. Misalignments, poor segmentation, and misplaced traces become more evident when objects are rendered in 3D, and annotation quality is improved when users switch between 2D and 3D views while working. PyReconstruct’s 3D scene includes several features to facilitate these actions. Double-clicking on a point in the 3D scene focuses on that point in the main window, providing a rapid way to identify 2D locations from the 3D space (Fig. 9B). Objects rendered in the 3D scene can be translated and rotated, allowing the user to customize the 3D view, for example, to visualize all objects of interest in a single glance (Fig. 9C). Objects from other series can also be imported into the current 3D scene. Customized 3D views can be saved as “scenes” that store object attribute and location information, which can be opened across annotation sessions. Scenes, in addition to aiding users in annotation, mitigate the need to export meshes to external 3D modeling software when making simple figures for publications.

### Under the hood

#### Interface framework, data structures, and legacy compatibility

PyReconstruct’s user interface is powered by Python bindings for Qt6, an opensource and platform-independent framework for developing graphical user interfaces (62). Users can therefore run PyReconstruct on most major modern operating systems (Windows, macOS, and many Linux distros) right out of the box.

Annotation data in legacy Reconstruct was stored over multiple “trace files”, one for each section, making moving series data between computers and users cumbersome and prone to data loss. Storing annotation data in a single, locally stored file facilitates data sharing amongst users. Annotation data in PyReconstruct is collected and stored in a single JSON-structured file with the extension .jser (portmanteau of “JSON” and “series”, pronounced “<u>jay</u>-sir”, Fig. 10). Image data are stored separately in a format and location defined by the user.

**Figure 10.**
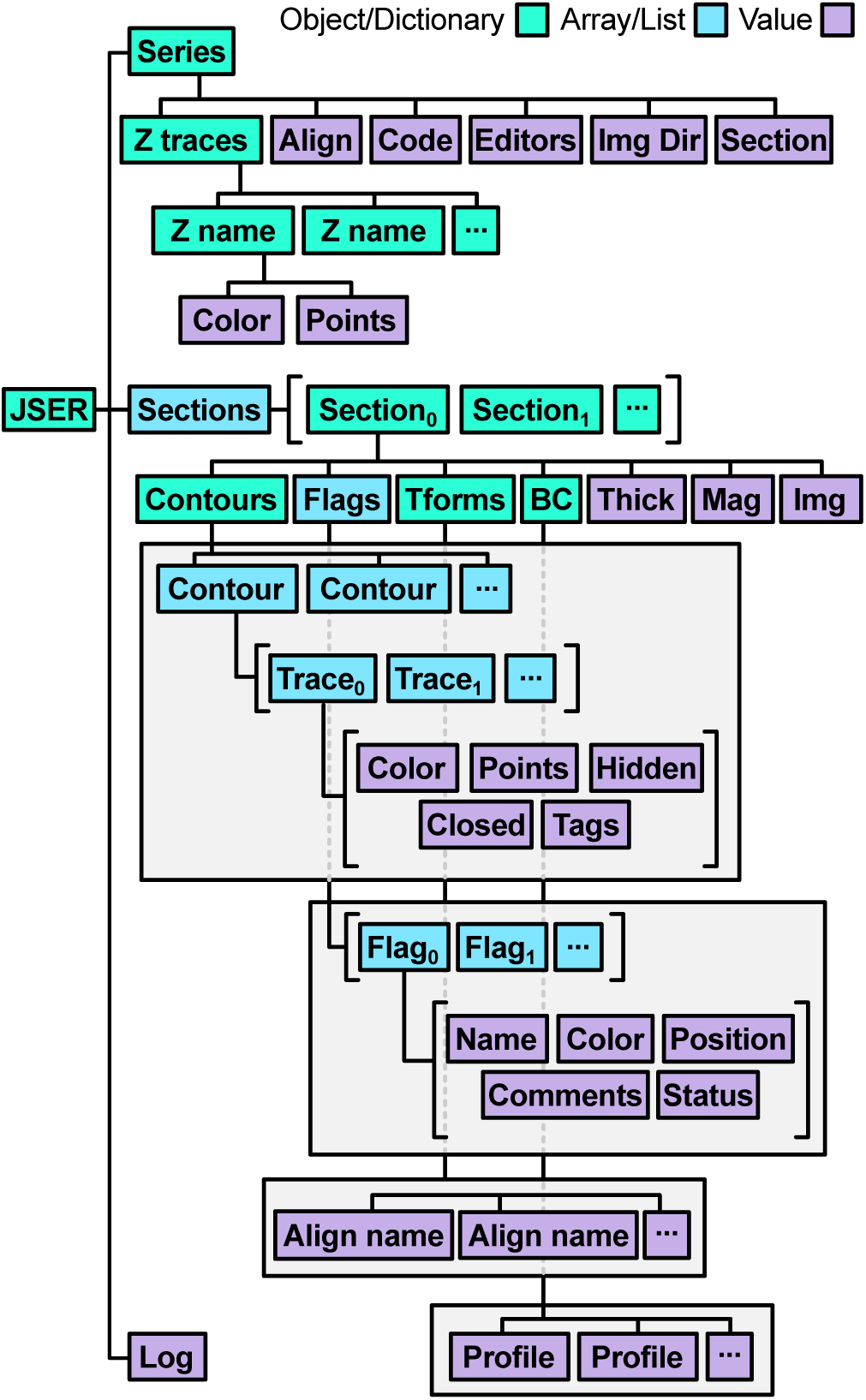
Schematic of PyReconstruct annotation data. Whereas legacy Reconstruct stored annotation data as multiple, non-canonical XML files, PyReconstruct stores annotation data locally as a single JSON-structured file (suffixed with .jser) loaded as a Python dictionary and edited using built-in modules or through PyReconstruct’s API. Series-wide data is held in a “Series” key, section data in a “Section key”, and series history in a “Log” key. The top-level key “Sections” maps to a list of section dictionaries with keys representing contour, flag, transformation data, etc. “Contours” is associated with a list of contours, which hold trace information. Traces are lists of trace attributes, such as trace color, points, visibility, etc. (Abbreviations: *Align* alignment, *Code* series code, *Img Dir* image directory, *Tforms* transforms, *BC* brightness/contrast profiles, *Thick* thickness, *Mag* pixel magnification, *Img* image file path.)

The jser file is divided into data pertaining to individual sections (section-specific data) and data pertaining to the entire series (series-specific data). Section-specific data contain contour information and annotations, transformations, and the section’s image attributes (magnification, brightness, contrast, and image file pointer). Series-specific data store information that spans multiple sections, including, for example, series meta-data, object attributes, current alignment, user information, and user preferences. All annotation data is accessible graphically through the application and through PyReconstruct’s Python API.

PyReconstruct implements a composite file pattern approach when loading and saving data: Data stored in the single jser file is decomposed at startup into discrete temporary hidden files that hold information pertaining to individual sections and the series. This approach optimizes RAM utilization while processing contour data and provides fault tolerance should the program unexpectedly close, or the user’s system fails.

Labs from a variety of fields continue to employ legacy Reconstruct as the primary tool used to annotate serial section EM data manually due in part to its ease of use. We therefore sought to make porting data from legacy Reconstruct into PyReconstruct simple. PyReconstruct is backward compatible with legacy structures: Series that were previously annotated in Reconstruct and saved as XML can be loaded into PyReconstruct by simply pointing to the legacy structures. No external conversion is necessary. Importantly, this process is bidirectional: Annotation data collected in PyReconstruct can also be exported as legacy XML structures in case users have established workflows that rely on legacy structures remaining intact, for example, with neuropil tools (36).

Features only compatible with PyReconstruct are saved in the jser file. Users therefore retain full access to the legacy application should they choose to do so and can still benefit from PyReconstruct’s modern annotation features.

#### Lifting RAM restrictions through multi-resolution scaling

Datasets loaded in legacy Reconstruct could exceed RAM; however, two sections were loaded into memory simultaneously. This meant large images needed to be cropped to a size compatible with the user’s RAM specs. Modern, ultra-large data viewers typically solve this issue by employing data structures that store chunked, compressed N-dimensional arrays at multiple resolution (or “scales”), such as with HDF5 or Zarr. Precomputed image datasets can be manipulated through a variety of wrappers such as h5py (63), PyTables (64), Z5 (65), and zarr-python (66). Instead of mapping entire images from disk into RAM, moving compressed chunks of data dramatically reduces loading times and makes navigating serial sections a much quicker and more pleasant experience. Saving the image data at multiple resolutions further reduces the amount of data loaded when users view low magnification views of a section.

To lift legacy Reconstruct’s RAM restrictions, we have implemented a multi-resolution Zarr approach to loading and viewing image data in PyReconstruct. Zarr was chosen as it was designed specifically for Python, supports robust multi-threading, provides access to customizable compressor and filter classes, and is used extensively in machine-assisted segmentation. Unlike applications that consolidate serial sections into a volumetric, 3D Zarr, PyReconstruct implements a 2D layer approach: each section being stored as a separate annotated group, which allows users to switch between the original images and Zarr at will.

Storing and retrieving image data in a chunk-wise manner means PyReconstruct users are in practice no longer RAM-limited when annotating series. Users are now virtually unbounded by series size, restricted only by the disk space available on their local machines, which can be augmented with external and remote storage. This strategy permits near-instantaneous loading of images in the field, as only chunks at the lowest possible resolution based on zoom are loaded during viewing. PyReconstruct maintains its flexible alignment strategy by applying affine transformations non-destructively, only after image data is loaded. A multi-resolution Zarr approach does increase the overall size of data stored per project, but PyReconstruct’s scaling strategy means Zarr datasets are never heavier than 1.33 times the size of the original image stack.

#### Interacting with external large-image applications and datasets

Volumetric datasets are much larger than they used to be, even recently (67–72). This is due in part to the transition from volumes constructed from small-field TEM images to those from high-throughput methods such as focused ion beam SEM (73, 74), serial block face SEM (75, 76), and tape and array methods (77–80). Relying on manual segmentation alone is therefore increasingly unfeasible. However, many labs (including our own) continue to employ manual segmentation as a primary technique. We therefore sought to provide a simple platform that annotators can use to begin to interact with the many automated segmentation tools being developed by the connectomics community.

Neuroglancer is a WebGL-based application for viewing, annotating, and querying large, multi-resolution volumetric data (81) and is the application of choice in several connectomics projects. Porting data to and from Neuroglancer offers one path through which automated segmentation efforts can be tapped into. PyReconstruct provides a graphical interface where users can export image data and contours as labeled Zarrs compatible with Neuroglancer. Data annotated in Neuroglancer can also be imported and converted to contour data for use in PyReconstruct.

## Discussion

PyReconstruct provides solutions that enhance manual segmentation of serial images in a user-friendly and familiar interface and offers several advantages over legacy Reconstruct. The target audience includes researchers from a variety of largely unrelated fields, who rely on manual annotating serial sections and volume EM data. Thus, PyReconsruct is a non-specialized tool that offers platform-independence, simplified data structures, a unique alignment strategy, and functionality to interact with a variety of external tools.

Integrated data quality control is often overlooked in annotation software, requiring users to develop their own external tracking systems. PyReconstruct has a robust and built-in curation system that addresses this critical gap. By embedding curation data directly within project files, PyReconstruct provides a streamed approach to team coordination and data integrity. PyReconstruct is fully opensource under a GNU General Public License v3 and users are encouraged to scrutinize and improve its source code, which is publicly available at GitHub (https://github.com/synapseweb/pyreconstruct). Opensource tools like PyReconstruct empower users to create custom solutions that can be integrated into the main product to the benefit the broader scientific community. This open environment also makes it possible to develop data conversion strategies that will allow PyReconstruct to accept annotations and segmentations from other systems.

As our understanding of alignment evolves, image registration strategies are advancing (46–57). The ability to re-align a series means users can adapt and refine their work at all stages, rather than being constrained to a single baked-in alignment. Image flaws resulting in alignment errors are often not noticed until the annotation stage, when users are viewing sections at higher magnifications. The ability to tweak existing alignments rapidly and create new alignments when required therefore facilitates annotation and improves the quality of the 3D outcomes. In PyReconstruct, users are at liberty to align the series, import alignments from external software, and store multiple alignments that can be switched to and from on demand. Importantly, this strategy enables users to employ simple, non-elastic alignments in series that suffer from image artifacts that warp portions of a section (see Fig. 7). Multiple alignment stacks for a single series and the ability to realign on demand also provides ground truth for developing more sophisticated automatic alignment methods.

Simplicity and generalizability come with tradeoffs. Fully segmenting, annotating, and curating EM volumes is time-consuming. For example, the complete manual annotation of a single, roughly 180 μm ^3^ volume of hippocampal tissue—an exceedingly small volume by today’s connectomics standards—took a team of expert annotators more than a year to complete and curate using legacy Reconstruct (82). The process of scaling up is therefore impractical (if not impossible) when annotators rely on manual segmentation alone. The push by many teams to produce semi- and fully automated segmentation pipelines has led to the segmentation of breathtakingly large datasets (67–69, 71). Nevertheless, these pipelines remain largely relegated to specialist communities. Therefore, a need remains to adopt automated segmentation approaches in simple user interfaces for non-specialist users.

One key challenge in adopting automated segmentation routines lies in balancing technical sophistication with accessibility. Modern automated segmentation tools require computational infrastructure and expertise in programming. The benefits of expanding access to these tools are immense. Moving forward, the developers of PyReconstruct are evolving its API and plugin framework to support the embedding of customized automated segmentation features into the software. This modular approach will enable researchers to leverage cutting-edge routines without the need for deep technical expertise, bridging the gap between specialist and non-specialist communities.

The evolution of imaging modalities over the past several decades has led to microscopy datasets that are larger than they have ever been (67–72, 83). These massive datasets have predominantly required server-based storage. Annotation tools that have grown up around these projects have therefore prioritized this approach (4, 9, 10). At present, PyReconstruct is a fully standalone application without server-client dependencies. Its users are not dependent on an internet connection, server maintenance, or remote data stores. While this approach offers simplicity and autonomy, users may need to generate manageable subvolumes from their data, especially when working with extremely large datasets.

Previously, researchers faced the onerous task of manually cropping individual serial images. Today, the widespread adoption of chunk-wise data storage formats and the increasing availability of standardized, open-source databases for large image data (84–88) have significantly mitigated these limitations. Users can efficiently extract subvolumes from remotely stored datasets through Python libraries like CloudVolume, which offer APIs to access precomputed volumes (89). These tools can be readily integrated into a workflow to sample regions from larger volumes and use the suite of annotation, curation, quantification, and visualization features to produce high quality data output from PyReconstruct.

## Acknowledgements

We would like to thank Harris lab members, GitHub users, members of our regular PyReconstruct user meetings, and students in NEU466G at the University of Texas at Austin for beta testing PyReconstruct and for their suggestions. Their input was vital in shaping the interface. We also would like to thank James Carson at the Texas Advanced Computing Center (TACC) for his help in integrating PyReconstruct with TACC’s online access points and Vijay Venu Thiyagarajan for his suggestions regarding data structures.

